# Drug Repurposing Screen Identifies an HRI Activating Compound that Promotes Adaptive Mitochondrial Remodeling in MFN2-deficient Cells

**DOI:** 10.1101/2025.06.23.660251

**Authors:** Prerona Bora, Mashiat Zaman, Samantha Oviedo, Sergei Kutseikin, Nicole Madrazo, Prakhyat Mathur, Meera Pannikkat, Sophia Krasny, Alan Chu, Kristen A. Johnson, Danielle A. Grotjahn, Timothy E. Shutt, R. Luke Wiseman

**Affiliations:** Department of Molecular and Cellular Biology, The Scripps Research Institute, La Jolla, CA 92037; Department of Biochemistry and Molecular Biology, Cummings School of Medicine, University of Calgary, Calgary, Alberta, Canada; Department of Integrative Structural and Computation Biology, The Scripps Research Institute, La Jolla, CA 92037; Calibr-Skaggs Institute for Innovative Medicines, The Scripps Research Institute, La Jolla, CA 92037; Departments of Medical Genetics and Biochemistry & Molecular Biology, Cumming School of Medicine, Hotchkiss Brain Institute, Snyder Institute for Chronic Diseases, Alberta Children’s Hospital Research Institute, University of Calgary, Calgary, Alberta, Canada

**Author notes:** To whom correspondence should be addressed: R. Luke Wiseman, Timothy E. Shutt. These authors contributed equally (Prerona Bora, Mashiat Zaman).

## Abstract

Pathogenic variants in the mitochondrial outer membrane GTPase MFN2 cause the peripheral neuropathy Charcot-Marie-Tooth Type 2A (CMT2A). These mutations disrupt MFN2-dependent regulation of diverse aspects of mitochondrial biology including organelle morphology, motility, mitochondrial-endoplasmic reticulum (ER) contacts (MERCs), and respiratory chain activity. However, no therapies currently exist to mitigate the mitochondrial dysfunction linked to genetic deficiencies in MFN2. Herein, we performed a drug repurposing screen to identify compounds that selectively activate the integrated stress response (ISR) – the predominant stress-responsive signaling pathway responsible for regulating mitochondrial morphology and function. This screen identified the compounds parogrelil and MBX-2982 as potent and selective activators of the ISR through the OMA1-DELE1-HRI signaling axis. We show that treatment with these compounds promotes adaptive, ISR-dependent remodeling of mitochondrial morphology and protects mitochondria against genetic and chemical insults. Moreover, we show that pharmacologic ISR activation afforded by parogrelil restores mitochondrial tubular morphology, promotes mitochondrial motility, rescues MERCs, and enhances mitochondrial respiration in *MFN2*-deficient cells. These results demonstrate the potential for pharmacologic HRI activation as a viable strategy to mitigate mitochondrial dysfunction in CMT2A and other pathologies associated with MFN2 deficiency.

## INTRODUCTION

While originally defined for its role in promoting mitochondrial fusion, the outer mitochondrial membrane (OMM) GTPase MFN2 is now recognized to have additional functions, including the maintenance of mitochondrial-endoplasmic reticulum (ER) contacts (MERCs) and mediating mitochondrial motility.^1–4^ These primary functions of MFN2 can lead to secondary roles for this protein involved in regulating other aspects of mitochondrial biology including respiration.^5–10^ Consistent with the importance of MFN2 for mitochondrial function, >150 mutations localized throughout the MFN2 structure are causatively associated with the neurological disease Charcot-Marie-Tooth Type 2A (CMT2A), a genetic peripheral neuropathy involving axonal degeneration and dysfunction.^8,11^ The mechanistic basis by which MFN2 dysfunction leads to CMT2A is not fully understood^8^, although reduced fusion of the mitochondrial network^11–13^, decreased ER-mitochondrial contacts^1,14^, and impaired mitochondrial motility^11,15^ have all been implicated. With CMT2A accounting for 30-40% of all axonal cases of CMT2^16–18^, significant effort has been directed at developing therapeutic strategies to mitigate these pathologic mitochondrial dysfunctions associated with impaired MFN2 activity. These efforts include strategies to enhance activity of the alternative OMM GTPase MFN1^12,19,20^, inhibit pathogenic MFN2 alleles^21^, and promote MFN2 activity.^22–26^ However, while progress has been made, no therapeutic approaches are currently available to ameliorate the pathologic mitochondrial dysfunction implicated in MFN2-associated CMT2A pathogenesis.

Another potential strategy to mitigate mitochondrial dysfunction associated with MFN2 deficiency is to pharmacologically enhance the activity of adaptive stress-responsive signaling pathways that regulate mitochondrial morphology and function. Recently, the integrated stress response (ISR) has emerged as the predominant signaling pathway responsible for adapting mitochondrial biology in mammalian cells. The ISR comprises four stress-activated kinases – GCN2, HRI, PERK, and PKR – that are activated in response to diverse types of pathologic insults including amino acid deprivation, mitochondrial stress, ER stress and viral infection, respectively.^27,28^ Upon activation, these kinases selectively phosphorylate the α subunit of eukaryotic initiation factor 2 (eIF2α), resulting in both the attenuation of new protein synthesis and the activation of stress-responsive transcription factors such as ATF4.^27,28^ ISR-dependent transcriptional and translational signaling regulates many aspects of mitochondrial biology such as proteostasis, morphology, lipid synthesis, and respiratory chain activity.^29–36^ Notably, ISR-dependent translational attenuation promotes mitochondrial elongation during ER stress downstream of PERK through a mechanism involving inhibition of the pro-fission mitochondrial GTPase DRP1.^30,32^ This finding suggests that pharmacologically enhancing ISR signaling offers a potential opportunity to mitigate pathologic mitochondrial fragmentation and subsequent organelle dysfunctions linked to MFN2 deficiency.

Consistent with this idea, pharmacologic activation of ISR kinases has proven beneficial for mitigating mitochondrial fragmentation and dysfunction in cellular models of diverse diseases. For example, treatment with the glutamyl-prolyl tRNA synthetase inhibitor halofuginone – a compound that activates the ISR kinase GCN2^37^ – promotes adaptive mitochondrial elongation and restores mitochondrial morphology and respiratory chain activity in cells deficient in the ISR kinase PERK.^29,31^ Similarly, pharmacologic activation of ISR kinases such as PERK or GCN2 mitigates mitochondrial fragmentation induced by the calcium ionophore ionomycin.^29^ However, despite this promise, currently available strategies to activate the ISR are limited in their translational potential for diseases like CMT2A by factors including poor selectivity for the ISR, low potency, dosing limitations, and off-target activities that directly impact mitochondrial biology.^31^ For example, the compound BtdCPU was previously shown to activate the ISR kinase HRI by inducing mitochondrial dysfunction through a mechanism involving mitochondrial uncoupling, precluding its use for correcting mitochondrial dysfunction in MFN2-deficient cells.^31^ Further, other ISR activators including the GCN2 activator halofuginone or ATP competitive inhibitors of ISR kinases can also induce ISR-independent translational inhibition or inhibit ISR kinase activity, respectively, at certain concentrations.^38–40^ Thus, new compounds are required to access adaptive ISR-dependent mitochondrial remodeling to further probe the potential for pharmacologic ISR activation to mitigate mitochondrial dysfunction associated with MFN2 deficiency.

Here, we employed a drug-repurposing screen to identify highly selective ISR activators with improved potential for mitigating mitochondrial dysfunction induced by reduced MFN2 activity. We screened the ReFRAME drug repurposing library comprising compounds previously shown to have promising safety profiles in human trials to identify compounds that selectively activate the ISR.^41^ This approach identified two compounds, parogrelil and MBX-2982, which we show potently activate the ISR with transcriptome-wide selectivity. Intriguingly, these compounds both activate the ISR through the OMA1-DELE1-HRI signaling axis; however, they do not induce the mitochondrial dysfunction commonly associated with the activation of this pathway (e.g., uncoupling, organelle fragmentation). Instead, we show that parogrelil and MBX-2982 promote ISR-dependent mitochondrial elongation in mammalian cell culture models and reduce mitochondrial fragmentation induced by chemical or genetic insults. Finally, we show that pharmacologic ISR activation using the HRI activator parogrelil mitigates diverse aspects of mitochondrial dysfunction in MFN2-deficient cells including morphology, motility, ER-mitochondrial contacts, and respiration. Collectively, these results demonstrate the potential for pharmacologic HRI activation to promote adaptive mitochondrial remodeling and mitigate mitochondrial dysfunction in MFN2-deficient cells, establishing pharmacologic ISR activation as a potential strategy to mitigate mitochondrial dysfunction in CMT2A and other diseases linked to reduced MFN2 activity.

## RESULTS

### Screening of the ReFRAME library to identify ISR activating compounds

We screened the 15,480 molecule ReFRAME drug repurposing library^41^ to identify compounds that activate the ATF4-FLuc ISR reporter stably expressed in HEK293T cells (**Fig. 1A**).^42^ The ER stressor thapsigargin (Tg) was used as a positive control for this primary screen to demonstrate robustness of the screening assay (robust Z’ factor = 0.5; S/N 2.53). Our primary screen identified 153 compounds that showed ATF4-FLuc activation that was >90% that observed for Tg (**Table S1**). These compounds were then subjected to dose response screening to define their AC_50_ for ATF4-FLuc activation (robust Z’ for dose response screen = 0.47). This analysis identified 112 ‘hit’ compounds with a measurable AC_50_ less than 10 µM (**Table S1**). These hits included numerous compounds that activate the ISR through known mechanisms including proteasome inhibition (e.g., bortezomib) or mitochondrial stress (e.g., antimycin A), validating the efficacy of our assay (**Table S1**). To prioritize compounds for further study, we focused on compounds with: 1) high potency for ATF4-FLuc activation, 2) low toxicity in HEK293T cells, as measured in previous screens performed at TSRI-CALIBR, 3) compound structures suitable for continued development, and 4) effective safety profiles. From this prioritization, we selected 5 compounds for further study – ciclesonide, capadenoson, velsecorat, MBX-2982, and parogrelil (**Fig. 1B**, **Fig. S1A**). We repurchased these compounds and re-tested their ability to activate the ATF4-FLuc reporter. Repurchased ciclesonide, capadenoson, and velsecorat did not potently activate the ATF4-FLuc reporter (**Fig. S1B**). Further, these repurchased compounds did not induce expression of the ISR target genes *CHAC1* and *ASNS* in HEK293T cells (**Fig. S1C**). Thus, we excluded these compounds from further study. In contrast, MBX-2982 (MBX) and parogrelil (PGL) both activated the ATF4-FLuc reporter with similar potency to that observed in our original screen (**Fig. 1C**) and did not show cytotoxicity in HEK293T cells (**Fig. S1D**). Co-treatment with the ISR inhibitor ISRIB – a compound that desensitizes cells to ISR-dependent phosphorylation of eIF2α^43–46^ – blocked ATF4-FLuc activation induced by MBX and PGL (**Fig. 1D**). This demonstrated that these compounds activate the ATF4-FLuc ISR reporter through an ISR-dependent mechanism. We then confirmed that MBX and PGL induced expression of the ISR target genes *CHAC1* and *ASNS* and showed dose-dependent increases in the expression of the ISR-regulated transcription factor ATF4 in HEK293T cells (**Fig. 1E,F**). This work identified MBX and PGL as non-toxic ISR activating compounds that we prioritized for further study.

**Figure 1.**
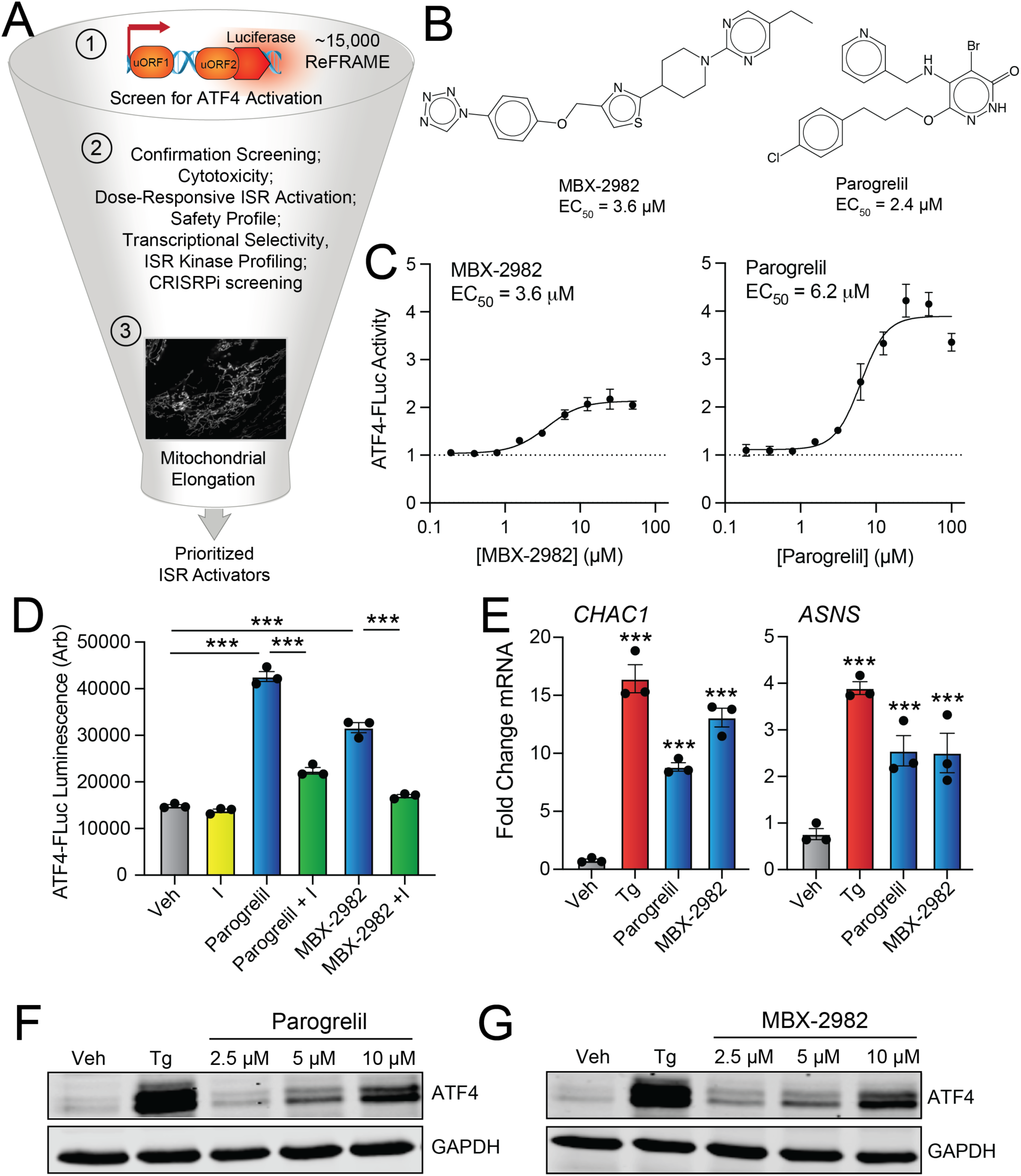
High throughput screening of the ReFRAME library identifies ISR activating compounds. **A.** HTS screening pipeline used to screen the ReFRAME library. **B**. Structures of the prioritized ISR activating compounds MBX-2982 and parogrelil. The AC_50_ for activation of the ATF4-FLuc ISR reporter determined from our primary screen is shown. **C**. ATF4-FLuc activation in HEK293T cells treated for 8 h with the indicated concentration of MBX-2982 (left) or parogrelil (right). **D**. ATF4-FLuc activation in HEK293T cells treated for 8 h with parogrelil (10 µM), MBX-2982 (10 µM), and/or ISRIB (200 nM). Error bars show SEM for n=3 replicates. **E**. Expression, measured by qPCR, of the ISR target genes *CHAC1* or *ASNS* in HEK293T cells treated for 6 h with thapsigargin (Tg; 500 nM), MBX-2982 (10 µM), or parogrelil (10 µM). Error bars show SEM for n=3 replicates. **F,G**. Immunoblots of lysates prepared from HEK293T cell treated for 6 h with thapsigargin (Tg; 500 nM) or the indicated dose of parogrelil (**F**) or MBX-2982 (**G**). ***p<0.005 for one-way ANOVA. Panel E statistics are shown relative to vehicle-treated cells.

### MBX-2982 and parogrelil are highly-selective activators of ISR signaling

We next defined the selectivity of ISR activation afforded by MBX and PGL. We performed RNAseq in HEK293T cells treated for 6 h with these compounds (**Table S2**). Intriguingly, the genes most significantly induced by these compounds were known transcriptional targets of the ISR (e.g., *CHAC, DDIT4, STC1*; **Fig. 2A,B**). Consistent with this observation, we found that treatment with MBX and PGL increased expression of a geneset comprising target genes regulated by the ISR (**Fig. 2C,D**; **Table S3**).^47^ However, we did not see significant increases in genesets comprising target genes of other stress-responsive signaling pathways such as the ATF and IRE1/XBP1s arms of the unfolded protein response (UPR), heat shock response (HSR), oxidative stress response (OSR), or hypoxic stress response.^47^ Similar results were observed when monitoring the expression of UPR, HSR, and OSR target genes by qPCR (**Fig. S2A,B**). These results indicate that MBX and PGL preferentially activate the ISR. Further supporting this idea, geneset enrichment analysis (GSEA) showed increased expression of UPR and mTOR genesets (**Fig. 2E,F**) – two genesets that include many ISR target genes. Similar increases in these genesets were observed for the ER stressor and potent activator of PERK-regulated ISR signaling thapsigargin (Tg; **Fig. S2C,D**). However, MBX also induced expression of the cholesterol biosynthesis geneset and suppressed expression of MYC target genesets, suggesting that this compound may have some additional activity. Collectively, these results indicate that MBX and, to a greater extent, PGL show transcriptome-wide selectivity for the ISR, particularly among stress-responsive signaling pathways.

**Figure 2.**
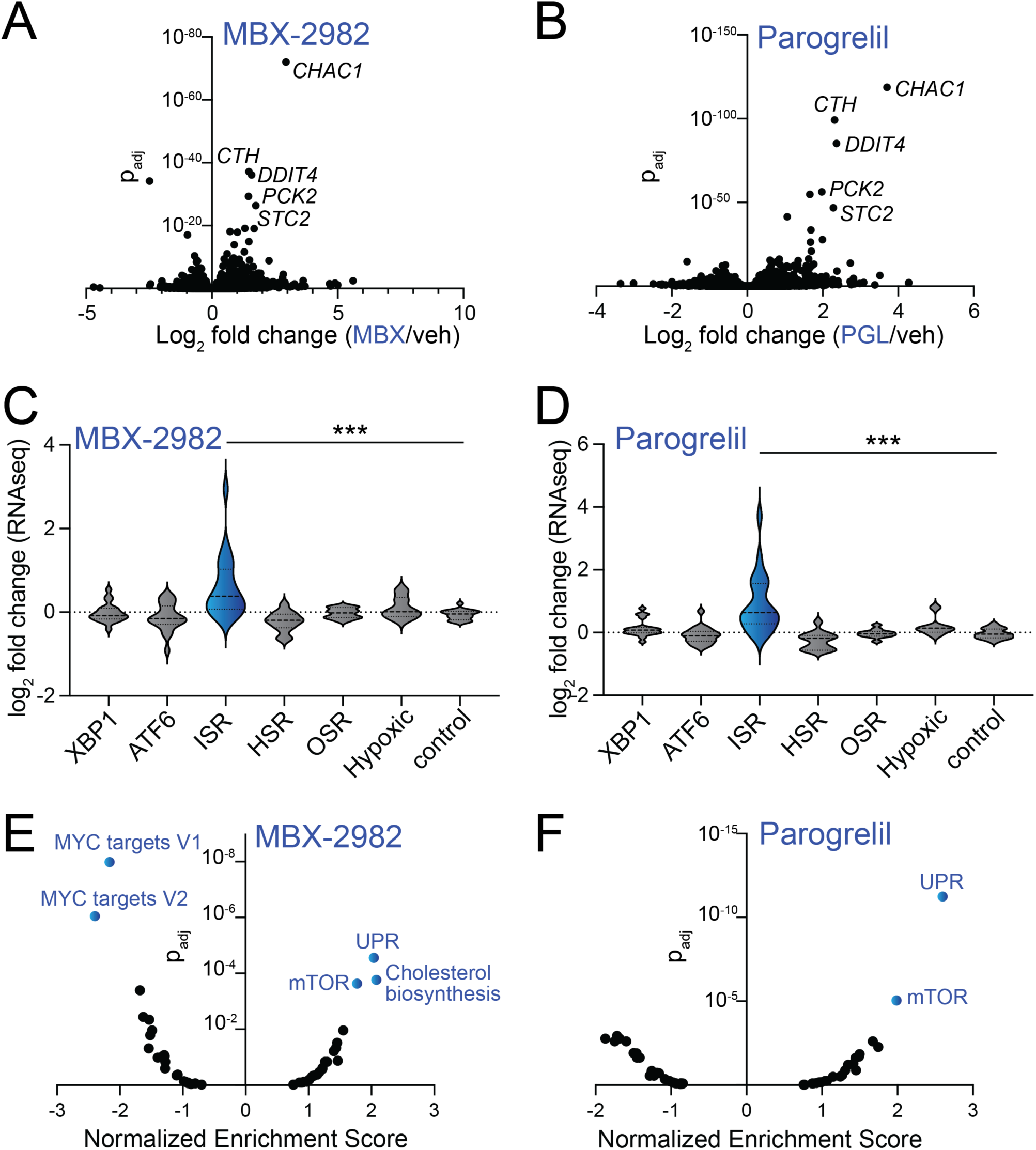
MBX-2982 and parogrelil selectively activate the ISR. **A,B.** Log_2_ fold change vs adjusted p-value (p_adj_) of gene expression, measured by RNAseq, in HEK293T cells treated for 6 h with MBX-2982 (**A**) or parogrelil (**B**). All RNAseq data is shown in **Supplemental Table S2**. **C,D**. Expression, measured by RNAseq of genesets comprising target genes of the XBP1s or ATF6 arms of the UPR, the ISR, HSR, OSR, or hypoxic stress response in HEK293T cells treated for 6 h with MBX-2982 (10 µM) or parogrelil (10 µM). A geneset comprising control genes is shown as a control. Genesets are described in **Supplemental Table S3**. ***p<0.005 for one-way ANOVA. **E,F.** Normalized enrichment score vs adjusted p-value (p_adj_) for geneset activation, measured by GSEA, in HEK293T cells treated for 6 h with MBX-2982 (10 µM; **E**) or parogrelil (10 µM; **F**). Full GSEA analysis is shown in **Supplementary Table S4.**

### MBX and PGL activate ISR signaling through the OMA1-DELE1-HRI signaling axis

We next wanted to identify the specific ISR kinase responsible for ISR activation afforded by treatment with MBX or PGL. We treated HEK293T cells stably expressing ATF4-mAPPLE CRISPRi depleted of individual ISR kinases with these compounds and monitored ISR activation by mAPPLE fluorescence.^48^ CRISPRi-depletion of *HRI*, but no other ISR kinases, blocked ISR activation induced by treatment with either MBX or PGL (**Fig. 3A,B**). Similarly, *HRI* depletion blocked expression of ISR target genes (e.g., *CHAC1*, *ASNS*) and increases of ATF4 protein in HEK293T cells treated with these two compounds (**Figs. 3C,D**). These results indicate that MBX and PGL activate the ISR kinase HRI.

**Figure 3.**
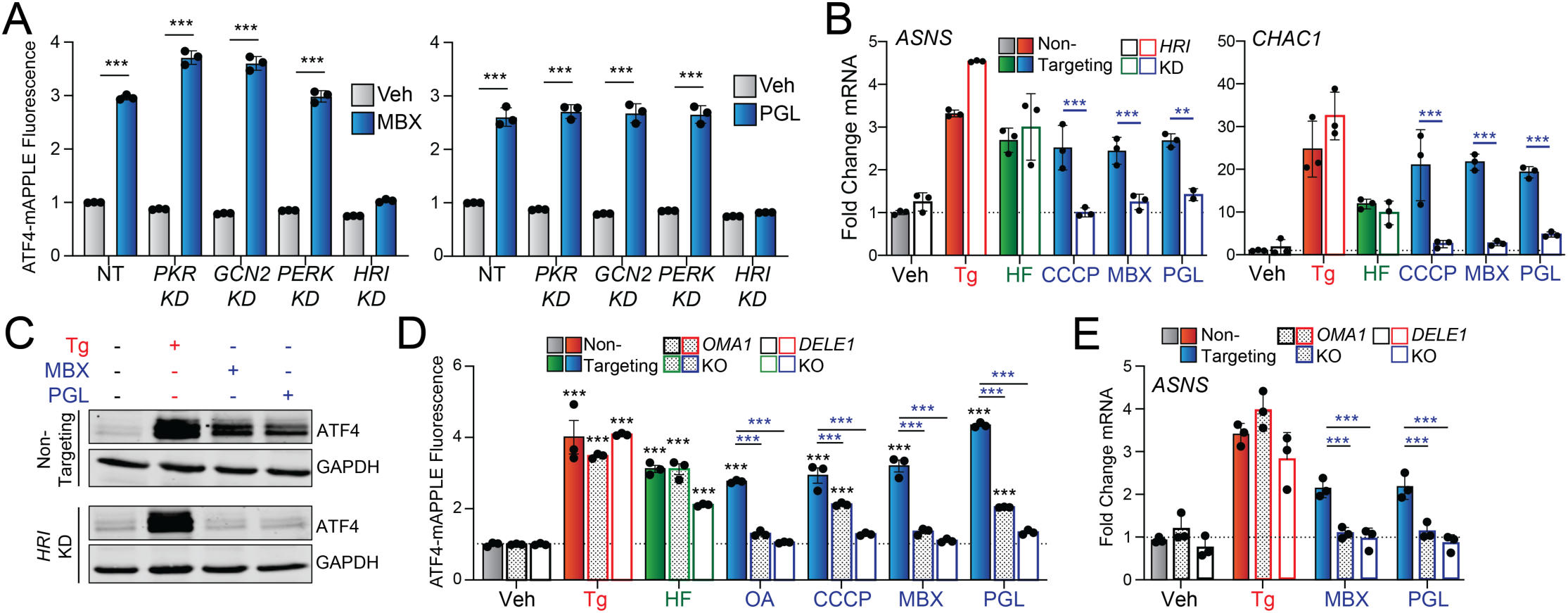
MBX-2982 and parogrelil activate the ISR through the OMA1-DELE1-HRI signaling axis. **A.** Activation of the ATF4-mAPPLE ISR reporter in HEK293T cells CRISPRi-depleted of the indicated ISR kinase treated for 16 h with MBX-2982 (MBX; 10 µM; left) or parogrelil (PGL; 10 µM; right). **B**. Expression, measured by qPCR, of the ISR target genes *ASNS* and *CHAC1* in non-targeting HEK293T cells or HEK293T cells CRISPRi-depleted *HRI* treated for 6 h with thapsigargin (Tg; 500 nM), halofuginone (100 nM), CCCP (10 µM), MBX-2982 (MBX; 10 µM) or parogrelil (PGL; 10 µM). **C**. Immunoblot of lysates prepared from non-targeting HEK293T cells (top) or HEK293T cells CRISPRi-depleted *HRI* (bottom) treated for 6 h with thapsigargin (Tg; 500 nM), MBX-2982 (MBX; 10 µM) or parogrelil (PGL; 10 µM). **D**. Activation of the ATF4-mAPPLE ISR reporter in HEK293T cells CRISPRi-depleted of *OMA1* or *DELE1* treated for 16 h with thapsigargin (Tg; 500 nM), halofuginone (100 nM), oligomycin A (OA; 500 nM), CCCP (10 µM), MBX-2982 (MBX; 10 µM) or parogrelil (PGL; 10 µM). **E**. Expression, measured by qPCR, of the ISR target gene *ASNS* in non-targeting HEK293T cells or HEK293T cells CRISPRi-depleted *OMA1* or *DELE1* treated for 6 h with thapsigargin (Tg; 500 nM), MBX-2982 (MBX; 10 µM) or parogrelil (PGL; 10 µM). Error bars show SEM for n=3 replicates. ***p<0.005 for one-way ANOVA (panel **A**) or two-way ANOVA (panels **B,D,** and **E**). Black asterisks show comparison with vehicle-treated controls. Colored asterisks show comparison for a given treatment across genotypes.

HRI is activated by diverse mechanisms including reductions in intracellular heme and proteolytic processing of the mitochondrial-targeted HRI adaptor protein DELE1 by the mitochondrial inner membrane protease OMA1.^42,48–50^ Thus, we sought to determine the dependence of MBX- and PGL-dependent HRI activation on these two mechanisms. We initially monitored ATF4-FLuc activation in HEK293T cells co-treated with the heme analog hemin (to reverse potential effects of heme depletion) and either MBX or PGL. Co-treatment with hemin did not impair the ability for these compounds to activate this ISR reporter (**Fig. S3A**). This result suggest that these compounds do not influence HRI activation through a mechanism involving reductions in intracellular heme. Next, we probed the impact of CRISPR deletion of *OMA1* or *DELE1* on ATF4-mAPPLE activation in HEK293T cells treated with MBX or PGL. Genetic depletion of *OMA1* partially inhibited ATF4-mAPPLE activation induced by MBX or PGL, while *DELE1* depletion completely inhibited ATF4-mAPPLE activation afforded by these compounds (**Fig. 3D**). Similar results were observed for oligomycin and CCCP (**Fig. 3D**) – two compounds previously shown to activate OMA1-DELE1-HRI signaling.^48,50^ In contrast, depletion of *OMA1* or *DELE1* did not significantly impact ATF4-mAPPLE activation afforded by treatment with the ER stressor thapsigargin (Tg) or the GCN2 activator halofuginone (HF), although we did observe a modest reduction in HF-dependent ATF4-mAPPLE fluorescence in *DELE1*-depleted cells. We further confirmed that *DELE1* or *OMA1* deletion impaired expression of ISR target genes (e.g., *ASNS*, *CHAC1*) and increases in ATF4 protein expression in HEK293T cells treated with MBX or PGL (**Fig. 3E**, **Fig. S3B,C**). These results indicate that MBX and PGL activate the ISR kinase HRI through the OMA1-DELE1-HRI signaling axis.

### MBX-2982 and parogrelil rescue mitochondrial fragmentation induced by pharmacologic or genetic insults

Activation of OMA1-DELE1-HRI signaling can be mediated by mitochondrial stressors such as the uncoupler CCCP and the ATP synthase inhibitor oligomycin, both of which also induce mitochondrial fragmentation by altering mitochondrial membrane potential.^48,50^ However, neither MBX nor PGL significantly impacted mitochondrial membrane potential in HEK293T cells, as measured by TMRE staining (**Fig. S4A**). Further, treatment with these compounds did not promote mitochondrial fragmentation in MEF cells (**Fig. 4A-C**). Instead, both MBX and PGL increased mitochondrial length and reduced organelle sphericity in these cells, indicating that pharmacologic HRI activation promotes mitochondria elongation – an adaptive phenotype previously shown to be induced by activation of other ISR kinases.^29,30,32,51^ Co-treatment with the ISR inhibitor ISRIB blocked mitochondrial elongation induced by these compounds (**Fig. 4A-C**). We confirmed that treatment with MBX or PGL induced expression of ISR target genes (e.g., *Asns*, *Chop*, *Chac1*) in MEFs by qPCR (**Fig. S4B**). Collectively, these results show that MBX and PGL can promote OMA1-DELE1-HRI signaling independent of mitochondrial uncoupling or mitochondrial fragmentation.

**Figure 4.**
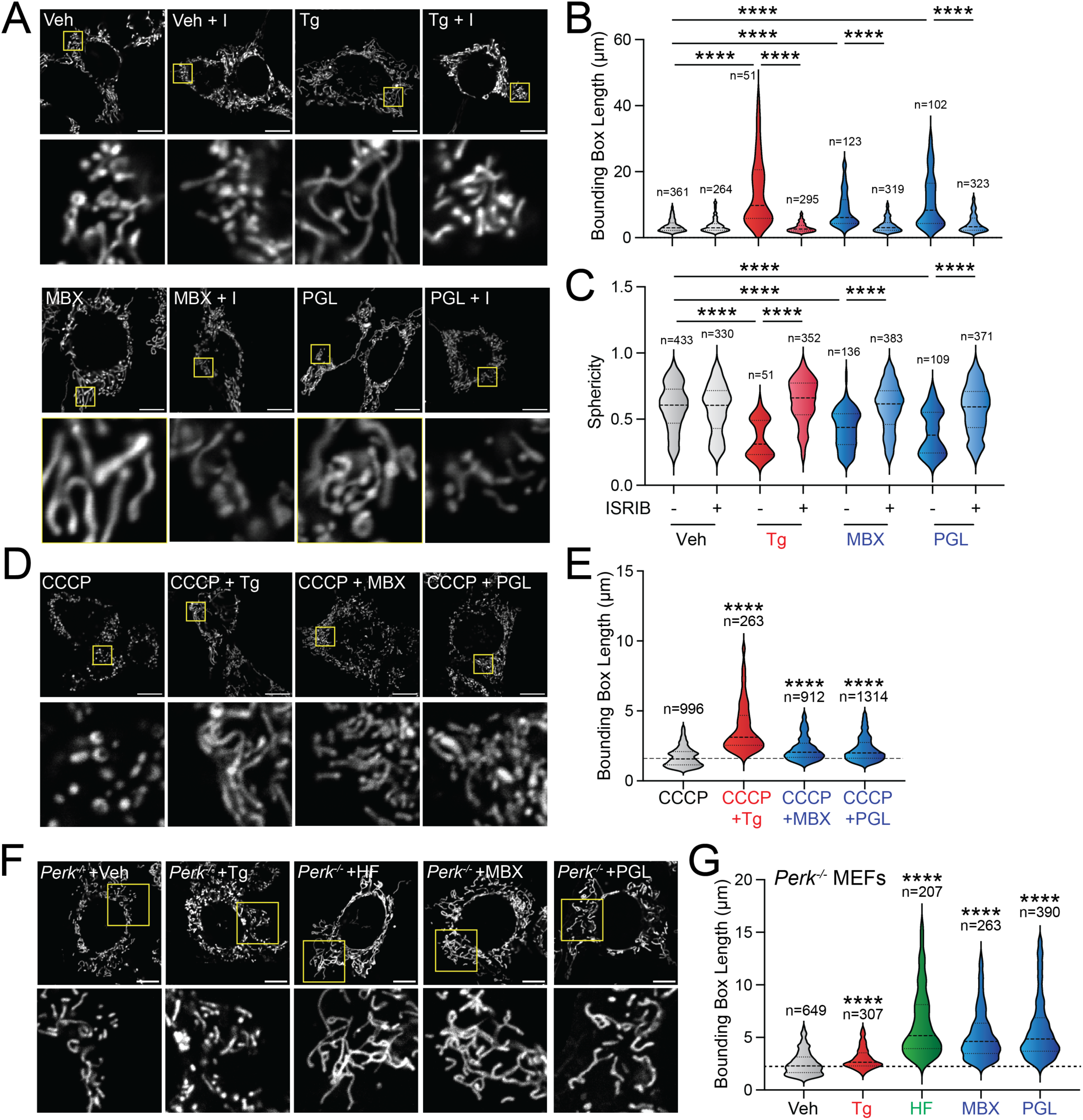
MBX and PGL promote protective mitochondrial elongation. **A.** Representative images of MEF cells stably expressing ^mt^GFP (MEF^mtGFP^) treated for 6 h with veh, thapsigargin (Tg; 500 nM), MBX-2982 (MBX; 10 µM), Parogrelil (PGL; 10 µM) with or without ISRIB (200 nM). The inset shows a 3-fold magnification of the region shown by the yellow box. Scale bars 10 µm. **B,C.** Quantification of bounding box axis (**B**) and sphericity (**C**) from the entire data set of images shown in (**A**). **D.** Representative images of MEF cells stably expressing ^mt^GFP (MEF^mtGFP^) pre-treated for 30 min with CCCP (10 µM) followed by 3 h treatment Tg (500 nM), MBX-2982 (MBX; 10 µM), Parogrelil (PGL; 10 µM). The inset shows a 3-fold magnification of the region shown by the yellow box. Scale bars 10 µm. **E.** Quantification of bounding box axis from the entire dataset of images shown in **D**. **F.** Representative images of TMRE stained *Perk^−/−^* MEF cells treated for 6 h with veh, Tg (500 nM), MBX-2982 (MBX; 10 µM), Parogrelil (PGL; 10 µM). The inset shows a 3-fold magnification of the region shown by the yellow box. Scale bars 10 µm. **G.** Quantification of bounding box axis from the entire dataset of images shown in **F**. ****p<0.001 for Kruskal-Wallis test. In panels **E** and **G**, comparisons are shown relative to vehicle-treated cells.

Our ability to promote ISR-dependent mitochondrial elongation using MBX or PGL suggests that treatment with these compounds could mitigate pathologic mitochondrial fragmentation induced by chemical or genetic insults. To test this notion, we pre-treated MEF cells with mitochondrial uncoupler CCCP for 30 min to induce mitochondrial fragmentation then monitored mitochondrial morphology in these cells following 3 h treatment with MBX or PGL. The ER stressor and ISR activator thapsigargin (Tg) was used as a control. This experiment showed that treatment with either MBX or PGL reduced mitochondrial sphericity and increased mitochondrial length in CCCP-treated cells (**Fig. 4D,E**, **Fig. S4C**). This demonstrates that treatment with these compounds can reduce CCCP-induced mitochondrial fragmentation. Further, we showed that MBX or PGL both rescued mitochondrial tubular morphology in MEF cells lacking the ISR kinase PERK (**Fig. 4F,G**, **Fig. S4D-I**)^52^ – a genetic perturbation previously shown to increase mitochondrial fragmentation.^51^ Collectively, these results indicate that MBX and PGL can mitigate mitochondrial fragmentation induced by both chemical and genetic insults by promoting adaptive ISR-dependent mitochondrial elongation.

### Pharmacologic ISR activation promotes adaptive mitochondrial remodeling in MFN2-deficient cells

MFN2 deficiency leads to pathologic mitochondrial dysfunctions implicated in CMT2A including organelle fragmentation, reduced motility, and decreased mitochondrial-ER contacts (MERCs).^8^ We sought to define the potential for pharmacologic ISR activation with MBX or PGL to mitigate these phenotypes in *Mfn2* knockout MEFs^4^ and U2OS cells CRISPR-deleted of *MFN2*. We also treated these cells with the GCN2 activator HF as a control for ISR activation. Treatment with HF or PGL for 6 h restored mitochondrial morphology to near wild-type levels in both MFN2-deficient MEF and U2OS cells (**Fig. 5A-C**, **Fig. S5A-C**), mirroring the rescue of mitochondrial morphology observed in cells treated with other chemical or genetic insults (see **Fig. 4**). Co-treatment with ISRIB blocked compound-dependent mitochondrial elongation in MFN2-deficient cells, confirming that this effect can be attributed to ISR activity. In contrast, MBX only modestly increased mitochondrial length in *Mfn2*-deficient MEFs and increased mitochondrial fragmentation in *MFN2*-deficient U2OS cells (**Fig. 5A-C**). This latter effect appears to be dependent on MFN2 deficiency, as treatment with MBX induced mitochondria elongation in *MFN2*-deficient U2OS cells re-expressing MFN2^WT^ (**Fig. S5C**). This observation suggests that there is a synthetic interaction between MBX and MFN2 deficiency that precludes the ability for this compound to promote adaptive mitochondrial remodeling in MFN2-deficient cells. Thus, we excluded MBX from further studies in cells lacking MFN2. Regardless, our results show that pharmacologic ISR activation with the GCN2 activator HF or the HRI activator PGL restores mitochondrial morphology in cells deleted of MFN2.

**Figure 5.**
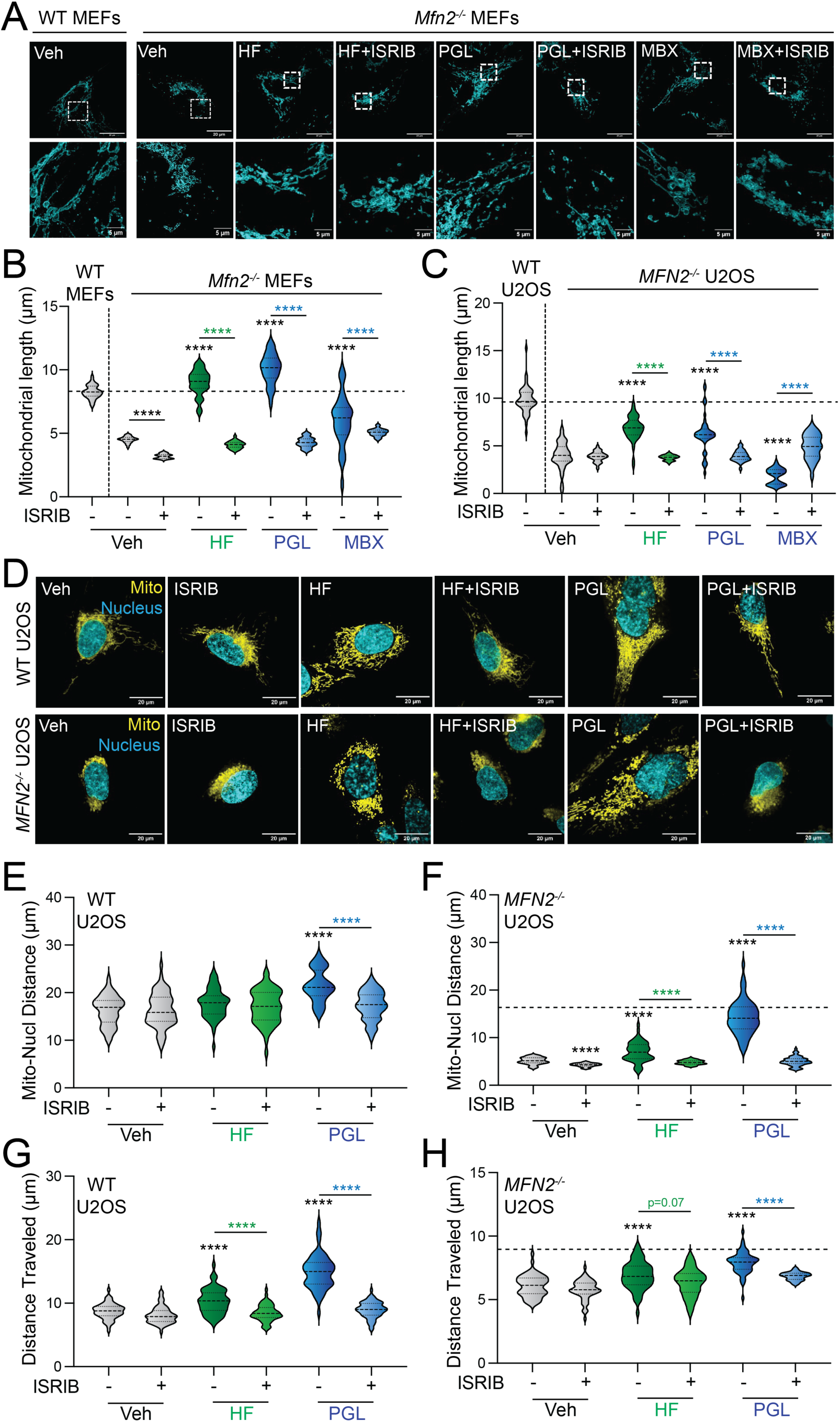
Pharmacologic ISR activation rescues mitochondrial morphology and motility in *MFN2*-deficient cells. **A**,**B**. Representative images (A) and quantification (B) of mitochondrial length in *Mfn2*-deficient MEFs treated for 6 h with halofuginone (HF; 100 nM); parogrelil (PGL; 10 µM), MBX-2982 (MBX; 10 µM), and/or ISRIB (200 nM) stained with TOMM20 antibodies (blue). Vehicle-treated wild-type (WT) MEF cells are shown as a control. **C**. Quantification of mitochondrial length in fixed *MFN2*-deficient U2OS treated for 6 h with halofuginone (HF; 100 nM); parogrelil (PGL; 10 µM), MBX-2982 (MBX; 10 µM), and/or ISRIB (200 nM) stained with TOMM20 antibodies. Vehicle-treated wild-type (WT) U2OS cells are shown as a control. Representative images are shown in **Fig. S5A**. **D**-**F**. Representative images (D) and quantification (E,F) of mitochondria-nuclear distance in wild-type U2OS cells or *MFN2-*deleted U2OS cells treated for 6 h with halofuginone (HF; 100 nM), parogrelil (PGL; 10 µM), and/or ISRIB (200 nM) stained with TOMM20 antibodies (yellow) and Hoechst (blue). The dashed line in (F) shows mitochondria-nuclear distance in vehicle-treated wild-type cells for comparison. **G**,**H**. Mitochondrial distance traveled, measured by time lapse live cell imaging, in wild-type (left) or *MFN2*-deficient U2OS cells (right) treated for 6 h with halofuginone (HF; 100 nM), parogrelil (PGL; 10 µM), and/or ISRIB (200 nM). The dashed line in (H) shows the distance traveled in vehicle-treated wild-type cells. Data shown were collected across two biological replicates. Violin plots show median and interquartile range. ****p<0.001 for Brown-Forsythe and Welch ANOVA. Black asterisks indicate comparison to vehicle-treated cells. Colored asterisks indicate comparison to cells co-treated with ISRIB.

MFN2-deficiency also increases perinuclear clustering of mitochondria (**Fig. 5D-F**; **Fig. S5D-F**). This observation is attributed to disruptions in mitochondrial trafficking (i.e., motility) in cells lacking MFN2.^4^ Intriguingly, treatment with the HRI activator PGL increased mitochondrial distribution in both wild-type and MFN2-deficient U2OS and MEF cells (**Fig. 5D-F**; **Fig. S5D-F**). This increase was blocked by co-treatment with ISRIB. However, treatment with the GCN2 activator HF only modestly increased mitochondrial distribution in U2OS cells and did not influence distribution in wild-type U2OS and MEF cells or in *Mfn2*-deficient MEFs. This finding suggested that this phenotype is selectively induced by pharmacologic HRI activators such as PGL. The increased mitochondrial distribution observed in PGL-treated wild-type and MFN2-deficient U2OS and MEF cells likely reflects increased mitochondrial motility. To test this idea, we used live cell time-lapse imaging to quantify the distance mitochondria travel in *MFN2*-deficient U2OS treated with ISR activators. This work showed that PGL treatment increased the distance mitochondrial travel in both wild-type and *MFN2-*deficient U2OS through an ISRIB-sensitive mechanism (**Fig. 5G,H**; **Supplemental Movie 1**). Similar results were observed in wild-type and *Mfn2* KO MEFs (**Fig. S5G,H**). HF treatment also modestly increased the distance mitochondrial traveled in wild-type and MFN2-deficient U2OS and MEF cells, albeit to a much lesser extent than that observed for PGL. Together, these results show that pharmacologic ISR activation, especially PGL-dependent HRI activation, restores organellar distribution and enhances mitochondrial motility in MFN2-deficient cells.

MFN2 facilitates formation and stability of mitochondrial-ER contacts (or MERCs).^1^ While there is debate over the specific role of MFN2 at these interorganellar contact sites^53–55^, it is established that deficiencies in MFN2 reduce MERCs.^56–58^ Consistent with this finding, we used an established proximity ligation assay to confirm reductions in the number and size of MERCs in MFN2-deficient MEF and U2OS cells (**Fig. 6A**, **Fig. S6A**).^10^ Treatment with the prioritized ISR activators HF or PGL increased the size, but not the number, of MERCs in wild-type U2OS cells (**Fig. S6B-D**). However, we found that these compounds restored both the size and number of MERCs in *MFN2*-deficient U2OS cells to levels similar or greater than those observed in wild-type cells (**Fig. 6B-D**). Co-treatment with ISRIB blocked these increases. Similar effects were observed in wild-type and *Mfn2*-deficient MEF cells treated with these ISR activators (**Fig. S6E-J**). These results show that pharmacologic ISR activation rescues MERCs in MFN2-deficient cells.

**Figure 6.**
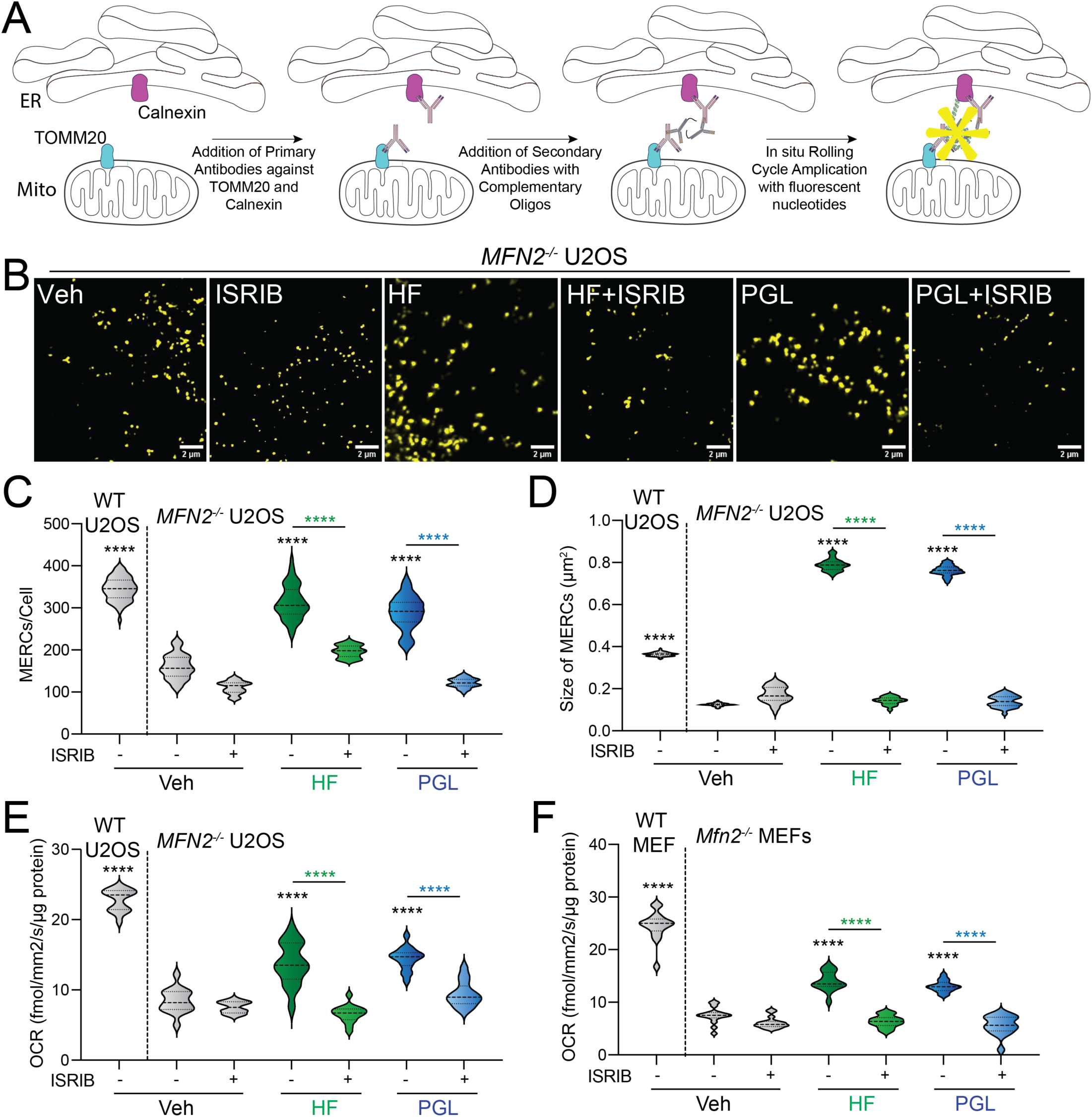
ISR activating compounds restore ER-mitochondrial contacts and enhance respiration in MFN2-deficient cells. **A.** Proximity ligation assay (PLA) used to monitor ER-mitochondrial contact sites. **B**-**D**. Representative images (**B**) and quantification of MERC number (**C**) and size (**D**), measured by PLA, in *MFN2*-deficint U2OS cells treated for 6 h with halofuginone (100 nM), parogrelil (PGL; 10 µM), and/or ISRIB (200 nM). PLA signal in wild-type U2OS cells is shown for comparison. **E,F**. Oxygen consumption rate (OCR), measured using the Resipher system, in MFN2-deficient U2OS (E) and MEF (F) cells treated for 6 h with halofuginone (HF; 100 nM), parogrelil (PGL, 10 µM), and/or ISRIB (200 nM). OCR in wild-type cells is shown as a control. ****p<0.005 for one-way ANOVA. Black asterisks show comparison with vehicle-treated cells and colored asterisks represents comparison with ISRIB co-treatment.

Apart from the CMT2A-associated phenotypes described above, deficiencies in MFN2 also reduces the basal oxygen consumption rate (OCR) in MEF and U2OS cells, as measured using the Resipher system (**Fig. 6E,F**). Thus, we sought to define the potential for pharmacologic ISR activation to restore respiratory activity in these cells. Treatment with PGL or HF increased OCR in wild-type cells through an ISRIB-sensitive mechanism (**Fig. S6I,J**). Similarly, we observed increased OCR in MFN2-deficient MEF and U2OS cells treated with these compounds (**Fig. 6E,F**). This increase was blocked by co-treatment with ISRIB. This finding indicates that, apart from morphologic aspects of mitochondrial biology, pharmacologic activation of HRI or GCN2 also enhances mitochondrial respiratory activity in MFN2-deficient cells.

## DISCUSSION

Here, we employed a drug repurposing screen to identify MBX-2982 (MBX) and parogrelil (PGL) as compounds that selectively activate the ISR through the OMA1-DELE1-HRI signaling pathway. We show that these compounds promote adaptive mitochondrial elongation that protects against mitochondrial fragmentation induced by chemical (e.g., CCCP) or genetic (e.g., *Perk* deficiency) insults. Further, we show that pharmacologic ISR activation afforded by the GCN2 activator HF or, especially, the HRI activator PGL ameliorates mitochondrial dysfunctions including fragmentation, reduced motility, decreased ER-mitochondrial contacts, and impaired respiration in MFN2-defiicent U2OS and MEF cells. These results establish MBX and PGL as highly selective ISR activating compounds and demonstrate their potential for correcting pathologic mitochondrial dysfunction in diseases associated with MFN2 deficiency such as CMT2A.

MBX and PGL are both previously identified drugs that successfully passed phase I clinical trials. MBX was originally described as a GPR119 agonist that was designed for the treatment of Type 2 diabetes^59^ , while PGL was identified as a PDE3 inhibitor that was developed for intermittent claudication.^60,61^ Herein, we show that these compounds are highly selective activators of the ISR. However, it is unlikely that either compound activates ISR signaling through these previously reported activities. GPR119 is not expressed in the cell lines used in this study, indicating that MBX-dependent ISR activation is independent of its role in targeting this G-protein coupled receptor. Further, the IC_50_ for PGL-dependent inhibition of PDE3, 0.26 nM, is significantly lower that observed for ISR activation, 5 µM.^61^ This suggests that PGL does not activate the ISR through PDE3. While efforts to identify the specific protein targets of these compounds required for ISR activation are currently ongoing, we show that both these compounds initiate ISR signaling through the mitochondrial stress-sensing OMA1-DELE1-HRI signaling axis. This finding is intriguing, as this signaling pathway had only previously been reported to be activated by mitochondrial stressors such as the uncoupler CCCP and the ATP synthase inhibitor oligomycin – treatments that also induce mitochondrial dysfunctions including organelle fragmentation, uncoupling, and impaired respiratory chain activity. In contrast, we show that activation of OMA1-DELE1-HRI signaling induced by MBX and PGL promotes adaptive ISR-dependent remodeling of mitochondria independent of these mitochondrial dysfunctions. This work highlights the potential for therapeutically accessing this endogenous mitochondrial stress-sensing mechanism using these compounds to promote mitochondrial remodeling and mitigate mitochondrial dysfunction in different diseases.

While ISR-dependent translational and transcriptional signaling induced downstream of the four different ISR kinases is generally considered to be similar, recent work has begun to suggest that HRI activation may have unique functions involved in adapting mitochondria during mitochondrial stress. Notably, HRI activation induced by iron chelation, but not stress-induced activation of other ISR kinases, was shown to promote PINK/PARKIN-independent mitophagy through a mechanism involving increased localization of phosphorylated eIF2α to the mitochondrial outer membrane.^62^ While our results show that pharmacologic activation of GCN2 with HF or HRI with PGL show similar improvements in mitochondria biology across most assays, we did note that PGL preferentially improved mitochondrial motility in MFN2-deficient U2OS and MEF cells, as compared to HF, through an ISRIB-sensitive mechanism. This observation suggests a previously unanticipated advantage for pharmacologically activating the HRI-regulated ISR, using compounds like PGL, for mitigating mitochondrial dysfunctions such as reduced motility associated with CMT2A and related disease.

Intriguingly, despite activating the ISR through OMA1-DELE1-HRI signaling and promoting adaptive mitochondrial elongation in wild-type cells, MBX demonstrated a synthetic interaction with MFN2 deficiency that limited its application for improving outcomes in cells lacking MFN2. This observation likely reflects an off-target activity of MBX, such as the MBX-dependent increase in cholesterol biosynthesis genes observed in our RNAseq analysis. Previous work has highlighted a role for MFN2 in regulating cholesterol, potentially explaining the synthetic interaction between MBX and MFN2 deficiency observed in our study.^63–66^ While MBX did not prove beneficial for correcting mitochondrial morphology in MFN2-deficient cells, this compound did ameliorate mitochondrial fragmentation induced by mitochondrial uncoupling or *Perk*-deficiency. This finding, combined with the selectivity of MBX for ISR activation, suggests that this compound could be useful for probing the potential for pharmacologic ISR activation to mitigate mitochondrial dysfunction induced by other types of pathologic insults.

No therapeutic approaches are currently available to mitigate pathologic mitochondrial dysfunction associated with MFN2 deficiency in CMT2A. Notably, PGL improved all three MFN2 functions implicated in CMT2A pathology: mitochondrial fragmentation, mitochondrial motility, and ER-mitochondrial contacts. Further, PGL enhanced respiratory chain activity in MFN2-deficient cells. Our results highlight the ability for pharmacologic ISR activation afforded by compounds such as the HRI activator PGL to inhibit mitochondrial fragmentation and dysfunction in MFN2-deficient cells. This finding suggests that treatment with these compounds could similarly ameliorate the myriad mitochondrial dysfunctions induced by hypomorphic CMT2A-associated MFN2 mutants that retain partial function.^8^ While most CMT2A MFN2 variants do not have deficient mitochondrial respiration, the fact that mitochondrial respiration was rescued in MFN2 deficient cells augurs well for the promise of ISR activation to rescue MFN2 dysfunction in CMT2A. Consistent with this notion, previous work has shown that HF-dependent ISR activation restores mitochondrial tubular morphology in patient fibroblasts expressing hypomorphic MFN2^D414V^.^29^ As we continue the development of this approach, it will be exciting to further probe the scope of CMT2A-associated MFN2 mutants that can be rescued by pharmacologic activation of HRI and other ISR kinases in patient-derived and animal models.

A challenge in translating pharmacologic ISR activators for diseases such as CMT2A is the potential for chronic activation of this pathway to lead to pathologic ISR signaling. However, pharmacologic ISR activation using compounds such as HF is well tolerated *in vivo*, highlighting the potential for the continued development of this approach.^67–69^ Further, both MBX and PGL have passed phase I clinical trials with good safety records, reflecting the *in vivo* potential for pharmacologic ISR activation using these compounds.^61,70^ As we continue developing pharmacologic ISR activators for CMT2A, and related diseases involving mitochondrial dysfunction, we will optimize pharmacokinetic and pharmacodynamic profiles of ISR activators to allow for the selective activation of adaptive, protective ISR signaling independent of chronic, pathologic ISR signaling, as we have for other stress pathway activators.^71,72^ Thus, as we advance the development of this class of compounds, we will be able to further establish pharmacologic ISR activation as a viable, broadly applicable strategy to mitigate mitochondrial dysfunction associated with etiologically diverse diseases.

## MATERIALS AND METHODS

### Plasmids, compounds, and reagents

All compounds used in this study were purchased: thapsigargin (Tg; 50-464-295, Fisher Scientific), ISRIB (SML0843, Sigma), oligomycin A (S1478, Selleck Chemicals), CCCP (C2759, Sigma), BtdCPU (32-489-210MG, Fisher), halofuginone (50-576-30001, Sigma), ciclesonide (S4646, Selleck Chemicals), capadenoson (HY14917, Med Chem Express), velsecorat (HY111453, Med Chem Express), MBX-2982 (1037792-44-1, Ambeed), and parogrelil (139145-27-0, Axon Medchem). Antibodies used were ATF4 (11815; Cell Signalling), Tubulin (T6074; Sigma), GAPDH (GTX627408; GeneTex), TOMM20, (ab186735; Abcam), and Calnexin (MAB3126; Sigma-Aldrich).

### Cell culture and transfections

HEK293T cells (purchased from ATCC), HEK293T cells stably expressing ATF4-FLuc or ATF4-mAPPLE CRISPRi-depleted of ISR kinases (*HRI*, *GCN2*, *PERK, PKR;* a kind gift from Martin Kampmann, UCSF)^42,48^ *and OMA1* or *DELE1, DELE1*-deficient HEK293T cells (kind gift from Xiaoyan Guo, Connecticut)^48^, and MEF^mtGFP^ cells (kind gift from Peter Schultz, TSRI)^73^ were all cultured at 37°C and 5% CO_2_ in DMEM (Corning-Cellgro) supplemented with 10% fetal bovine serum (FBS; GIBCO), 2 mM L-glutamine (GIBCO), 100 U/mL penicillin, and 100 mg/mL streptomycin (GIBCO). *Perk^+-+^* and *Perk^− / −^* MEFs (kind gifts from David Ron, Cambridge)^52^ were cultured at 37°C and 5% CO_2_ in DMEM (Corning-Cellgro) supplemented with 10% FBS (GIBCO), 2 mM L-glutamine (GIBCO), 100 Units/mL penicillin, 100 mg/mL streptomycin (GIBCO), non-essential amino acids (GIBCO), and 2-mercaptoethanol (ThermoFisher). MFN2 knockout MEFs (obtained from ATCC; CRL-2993)^4^ and MFN2 knockout U2OS (a generous gift from Dr. Edward A Fon, McGill University, Canada) were cultured at 37°C and 5% CO_2_ in DMEM high glucose media (Gibco), supplemented with 10% FBS.

### ReFRAME Screen

To complete the primary screen of the ReFRAME library, the ATF4-FLuc ISR reporter stably expressed in HEK293T cells were plated into 1536 well, with solid bottom Geiner plates with 750 cells per well in 5 µL of media. After allowing the cells to adhere overnight, the ReFRAME library compounds were transferred to the plates using Echo acoustic transfer to deliver 5 nL of compound per well, with a final screening concentration of 10 µM. After 24 hours, 5 µL of OneGlo (Promega) was added, the plates were centrifuged, incubated 10 minutes and quantified using the Luminescence module on the Pherastar (BMG). All hits which met the cut off criteria, were picked for hit confirmation in 10 point dose response curves, with a 1:3 dilution, beginning at 10 µM as the top dose in triplicate. All confirmed hits which had an EC_50_ < 10µM were evaluated in subsequent experiments.

### Measurements of ISR activation in ATF4-reporter cell lines

ATF4-FLuc reporter cells were seeded at a density of 15,000 cells per well in Greiner Bio-One CELLSTAR flat 384-well white plates with clear bottoms. The following day, cells were treated with the indicated compound in triplicate 10-point dose response format for eight hours. After treatment, an equal volume of Promega Bright-Glo substrate was added to the wells and allowed to incubate at room temperature for 10 minutes. Luminescence was then measured in an Infinite F200 PRO plate reader (Tecan) with an integration time of 1000 ms.

ATF4-mApple reporter cells were seeded at a density of 8,000 cells per well in 96-well TC-treated flat bottom plates (Genesee Scientific). The following day, cells were treated for 16 h with the indicated concentrations. Following treatment, cells were washed twice with phosphate-buffered saline (PBS) and dissociated using TrypLE Express (ThermoFisher). The enzymatic reaction was neutralized through addition of sorting buffer containing PBS and 5% FBS. Flow cytometry was performed on a Quanteon 4025 with NovoSampler Q. mApple (568/592 nm) was measured using the 561 nm green-yellow laser in combination with the 577/15 filter. Analysis was performed using FlowJo Software (BD Biosciences).

### Immunoblotting

Whole cells were lysed on ice in RIPA lysis buffer (50 mM Tris, pH 7.5, 150 mM NaCl, 0.1% SDS, 1% Triton X100, 0.5% deoxycholate and protease inhibitor cocktail (Roche)). Total protein concentrations of lysates were then normalized using the Bradford protein assay and lysates were combined with 1x Laemmli buffer supplemented with 100 mM DTT and boiled for 5 min. Samples (100 μg) were then separated by SDS-PAGE and transferred to nitrocellulose membranes (Bio-Rad). Membranes were blocked with 5% milk in tris-buffered saline (TBS) and then incubated overnight at 4 °C with the indicated primary antibody. The next day, membranes were washed in TBS supplemented with Tween, incubated with the species appropriate secondary antibody conjugated to IR-Dye (LICOR Biosciences), and then imaged using an Odyssey Infrared Imaging System (LICOR Biosciences). Primary antibodies were acquired from commercial sources and used in the indicated dilutions in antibody buffer (50 mM Tris [pH 7.5], 150 mM NaCl supplemented with 5% BSA (w/v) and 0.1% NaN3 (w/v)). ATF4 (1:500), Tubulin ( 1:5000), GAPDH (1:1000).

### Reverse Transcriptase-quantitative polymerase chain reaction (RT-qPCR)

The relative mRNA expression of target genes was measured using RT-qPCR. HEK293T cells were plated at 200,000 cells per well in a 6-well plate. After 24 h, the cells were treated with the indicated concentration of compound for 6 h. Cells were then washed with phosphate buffered saline (PBS; Gibco). RNA was extracted using Quick-RNA MiniPrepKit (Zymo Research) according to the manufacturers protocol. RNA (500 ng) was then converted to cDNA using the High Capacity Reverse Transcription Kit (Applied Biosystems). qPCR reactions were prepared using Power SYBR Green PCR Master Mix (Applied Biosystems), and primers (below) were obtained from Integrated DNA Technologies. Amplification reactions were run in an ABI 7900HT Fast Real Time PCR machine with an initial melting period of 95 °C for 5 min and then 45 cycles of 10 s at 95 °C, 30 s at 60 °C.

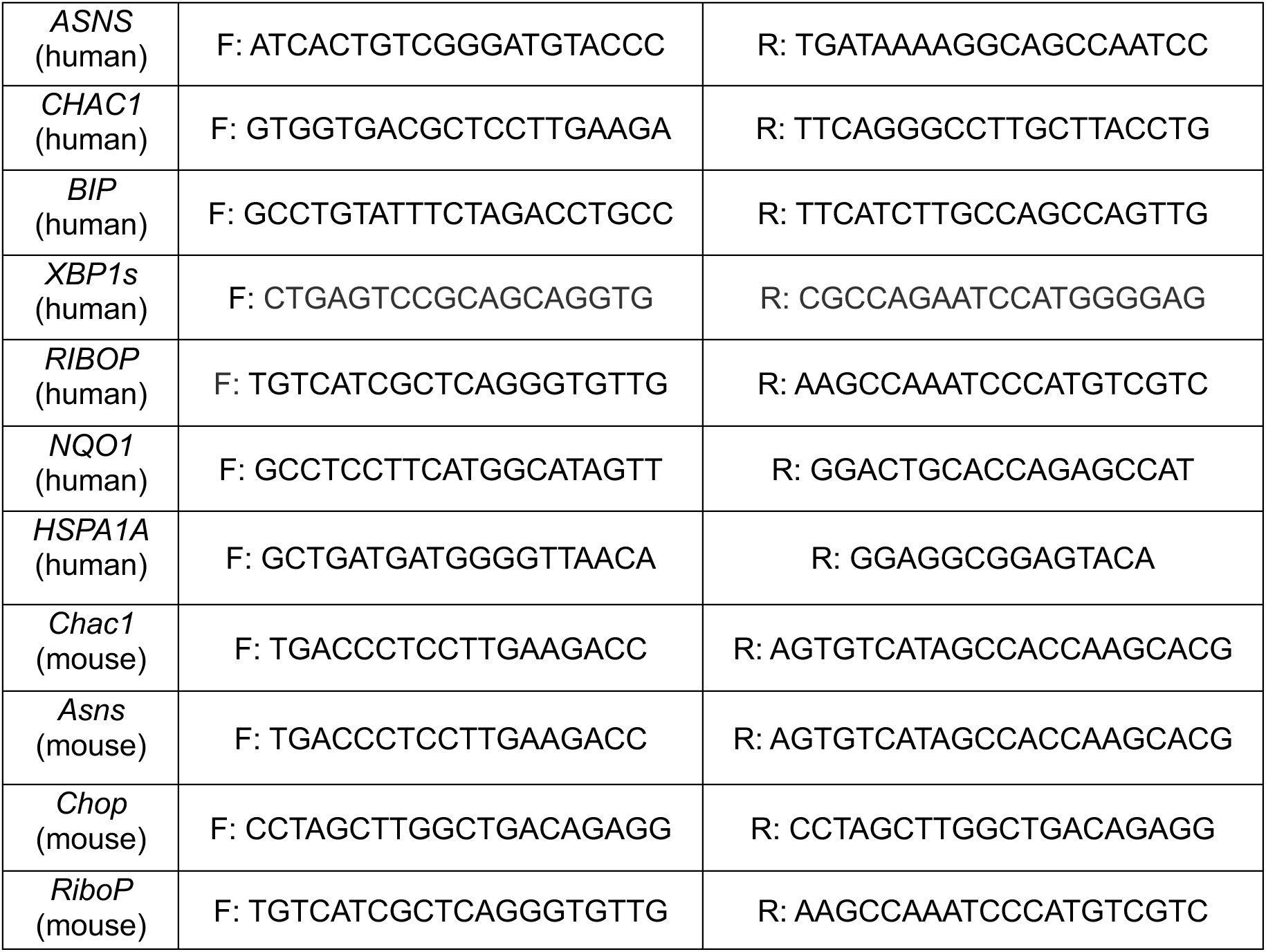

### RNA sequencing

HEK293T cells were plated at 200,000 cells per well in a 6-well plate. After 24 h, the cells were treated with the indicated concentration of compound for 6 h. Cells were then washed with phosphate buffered saline (PBS; Gibco). RNA was extracted using Quick-RNA MiniPrepKit (Zymo Research) according to the manufacturers protocol. RNA was submitted to TSRI genomics core for strand-specific mRNA sequencing on their DNBSEQ platform with three biological replicates per condition. All samples were sequenced to a minimum depth of 20 M PE 150 bp stranded reads. Alignment of reads was performed using DNAstar Lasergene SeqMan NGen 17 to the human genome GRCh37-dbSNP150 assembly. Aligned reads were imported into ArrayStar 17 as linear counts data. Differential expression analysis and statistical significance comparisons were assessed using DESeq 2 v. 1.34 in R compared to vehicle-treated cells. fGSEA (v1.20.0) was performed in R using the Hallmark Pathway Geneset (v7.5.1) from MSigDB. Code can be provided upon request.

### Fluorescence Microscopy

MEF^mtGFP^ and *Perk^+-+^* and *Perk^− / −^* MEFs were seeded at a density of 15,000 cells per well in eight-chamber slides (Ibidi) coated with poly-d-lysine (Sigma). The next day cells were treated with the indicated dose of compound for the indicated time. After treatment, cells were imaged on a Zeiss LSM 880 Confocal Laser Scanning Microscope equipped with a full incubation chamber for regulating temperature and CO_2_ during live-cell imaging. *Perk^+-+^* and *Perk^− / −^* MEFs were visualized by staining with TMRE (Thermofisher). Mitochondrial bounding box length and sphericity were quantified using the Imaris software, as previously described.^29^

Mitochondrial network morphology in wild-type and *Mfn2* KO MEFs and wild-type and *MFN2*-deficient U2OS cells was analyzed using immunofluorescence labelling. 20,000 cells were seeded in 12 mm coverslips and grown for 24 hours. Following compound treatment for 6 h, cells were subsequently fixed with 4% paraformaldehyde, permeabilized with 0.1% Triton X-100 and blocked with 10% FBS. The mitochondrial network was labelled using rabbit anti-TOMM20 (Abcam; ab186735) and visualized using anti-rabbit Alexa Fluor 488 (Thermo Fisher Scientific; A-11029). Coverslips were mounted using ProLong Diamond Anti-Fade mounting media (Invitrogen; P36965). Imaging was done using an Olympus Spinning Disk Confocal System (Olympus SD-IX83-SPINSR) using a 60× oil immersion objective. A cellVivo live cell incubator system was used for live cell imaging at 37°C and 5% CO_2_. Mitochondrial branch length was analyzed using the MiNA plugin^74^ on FIJI.^75^ Particle number and size were calculated using the Analyze Particles plugin on FIJI while Mito-nucleus distances were calculated as previously described for nucleus-lipid droplet distances.^9^

### Proximity ligation assay

Mito-ER contact sites were visualized using a previously established proximity ligation assay (PLA).^10^ Briefly, MEF and U2OS cells were seeded at a density of 30,000 cells per well in coverslips and grown for 24 hours. Following treatment with compounds for the indicated time, cells were fixed with PFA, permeabilized and blocked using the Duolink Blocking buffer from the PLA kit (DUI92008; Sigma-Aldrich). This step was followed by incubation with the primary antibodies (anti-TOMM20 for mitochondria and anti-Calnexin for the ER) overnight and subsequently with anti-mouse (DUO92004; Sigma-Aldrich) and anti-rabbit (DUO92002; Sigma-Aldrich) PLA probes. Manufacturer’s instructions were followed for addition of ligase and polymerase from the PLA kit. Secondary antibodies were added to visualize the mitochondria (AlexaFluor 488, A-11029; Thermo Fisher Scientific) and ER (AlexaFluor 647, A-21245; Thermo Fisher Scientific). The coverslips were mounted with ProLong Diamond Antifade Mountant before imaging.

### Oxygen Consumption Rate

Oxygen consumption rates were measured using the Lucid Resipher system. Wild-type and MFN2-deficient MEF and U2OS cells were seeded at a density of 15,000 cells per well in a 96-well plate and allowed to settle for 24 hours. Subsequently, drugs were added at the indicated concentrations and the lid was attached to the plate, which was attached to the hub. The oxygen consumption rate was measured at 6 hours and normalized to total protein present in the well. Total protein was calculated by collecting cells from each well, lysed using RIPA buffer (89901; Fisher Scientific) and quantified using the BCA assay, normalised to BSA.

### TMRE staining and flow cytometry

HEK293T cells were seeded at a density of 8,000 cells per well in 96-well TC-treated flat bottom plates (Genesee Scientific). The following day, cells were treated for 3 h with the compounds and the incubated with TMRE (200 nM) for 20 mins. Cells were washed twice with phosphate-buffered saline (PBS) and dissociated using TrypLE Express (ThermoFisher). The enzymatic reaction was neutralized through addition of sorting buffer containing PBS and 5% fetal bovine serum (FBS). Flow cytometry was performed on a Quanteon 4025 with NovoSampler Q. Fluorescence intensity of TMRE (552/574 nm) was measured using the 561 nm green-yellow laser in combination with the 577/15 filter. Analysis was performed using FlowJo Software (BD Biosciences).

### CellTiterGlo viability assays

HEK293T cells were seeded into flat black, poly-d-lysine coated 96-well TC-treated flat bottom plates (Genesee Scientific). The next day cells were treated with the indicated dose of compound for 24 h. After treatment, cells were lysed by the addition of CellTiter-Glo reagent (Promega). Samples were dark adapted for 10 min to stabilize signals. Luminescence was then measured in an Infinite F200 PRO plate reader (Tecan) and corrected for background signal. All measurements were performed in biologic triplicate.

## ACKNOWLEDGEMENTS

We thank Ilia Droujinine and Rama Aldakhlallah for comments on the manuscript, and the microscopy and genomic cores at TSRI for experimental support. We also thank the Wellcome Trust and the Bill & Melinda Gates Foundation for their support of the ReFRAME library. This work was supported by the National Institutes of Health (NS095892 and NS125674 to RLW), the Canadian Institutes of Health Research (to TES), George E. Hewitt Foundation Postdoctoral Fellowship (to PB), a Hotchkiss Brain Institute International Graduate Recruitment Scholarship (to MZ), a National Science Foundation Predoctoral Fellowship (to SO), and an American Heart Association Predoctoral Fellowship (to PM).

---

**Figure S1.**
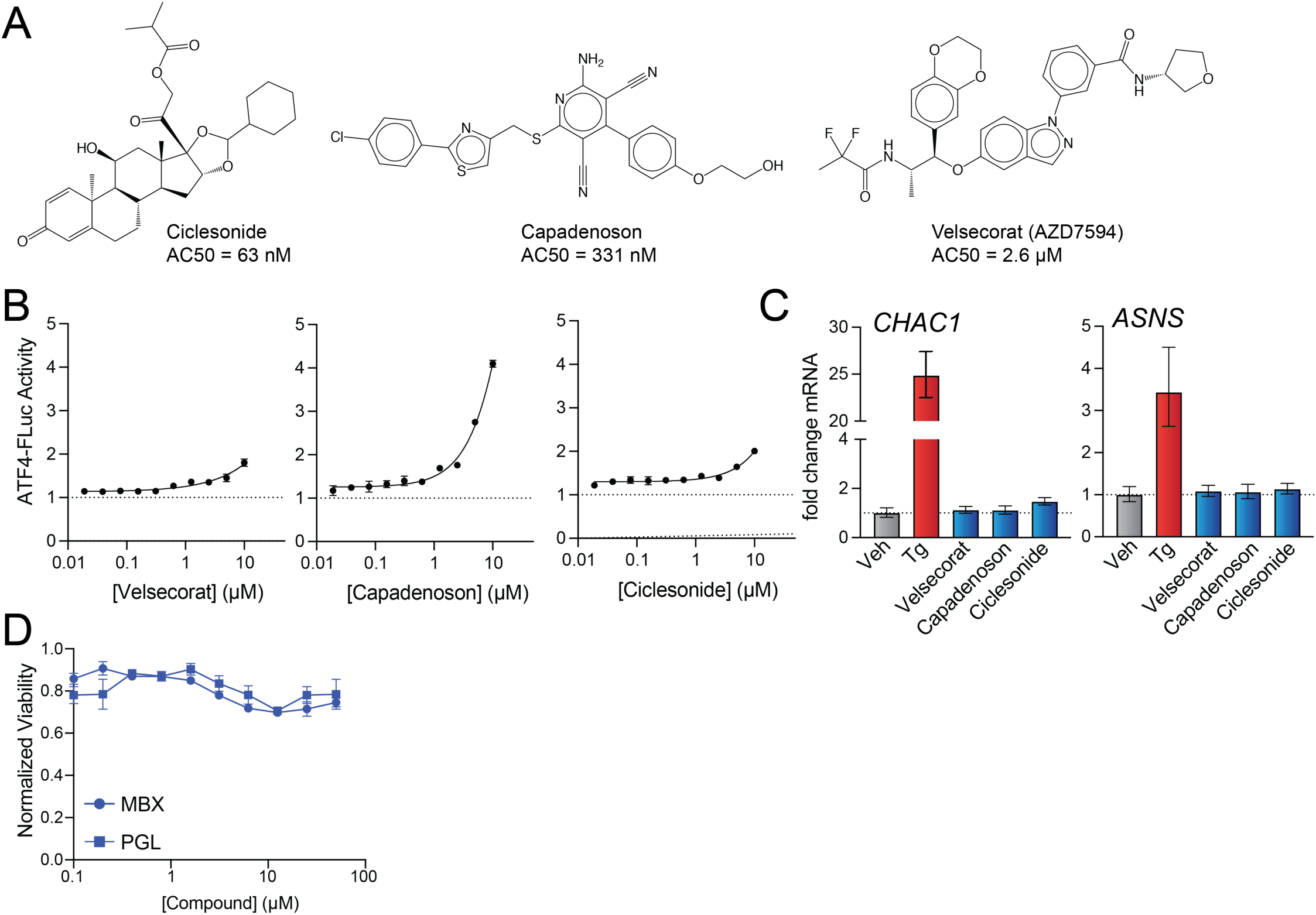
(Supplement to Figure 1). High throughput screening of the ReFRAME library identifies ISR activating compounds. **A**. Structures of ciclesonide, capadenoson, and velsecorat. **B**. ATF4-FLuc activation in HEK293T cells treated for 8 h with the indicated concentration of velsecorat (left), capadnoson (middle), and ciclesonide (right). Error bars show SEM for n=3 replicates. **C**. Expression, measured by qPCR, of the ISR target genes *CHAC1* or *ASNS* in HEK293T cells treated for 6 h with thapsigargin (Tg; 500 nM), velsecorat (10 µM), capadnoson (10 µM), or ciclesonide (10 µM). Error bars show SEM for n=3 replicates. **D**. Viability, measured by Cell Titer Glo, of HEK293T cells treated for 24 h with the indicated concentration of MBX-2982 (MBX) or parogrelil (PGL).

**Figure S2.**
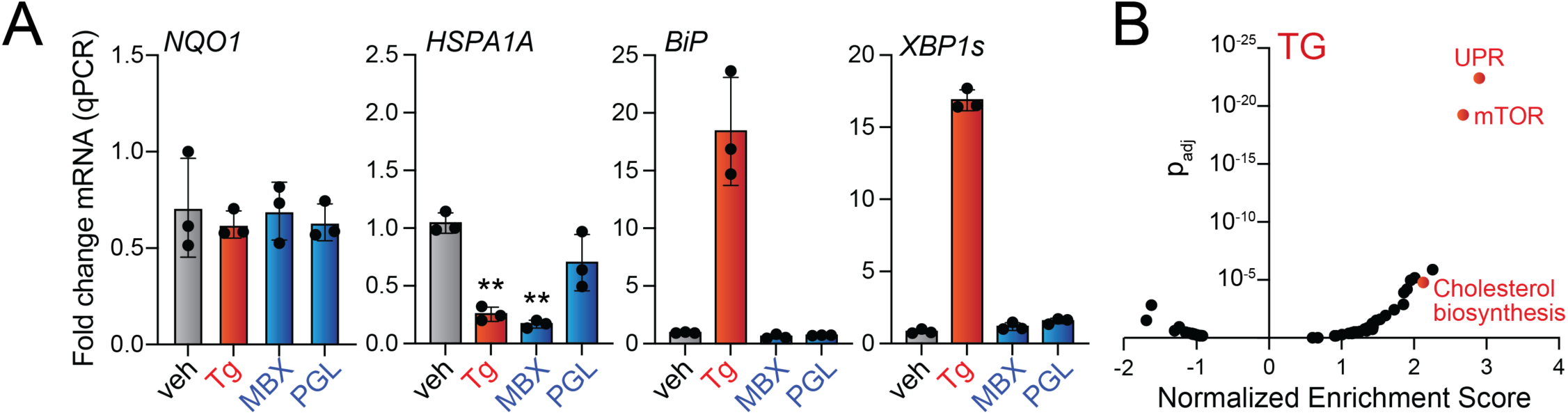
(Supplement to Figure 2). MBX-2982 and parogrelil selectively activate the ISR. **A.** Expression, measured by qPCR, of the OSR target gene *NQO1*, the HSR target gene *HSPA1A*, and the UPR target genes *BiP* or *XBP1s* in HEK293T cells treated for 6 h with thapsigargin (Tg; 500 nM) MBX-2982 (10 µM) or parogrelil (PGL; 10 µM). **B**. Normalized enrichment score vs adjusted p-value (p_adj_) for geneset activation, measured by GSEA, in HEK293T cells treated for 6 h with thapsigargin (Tg; 500 nM). Full GSEA analysis is shown in **Supplementary Table S4.**

**Figure S3.**
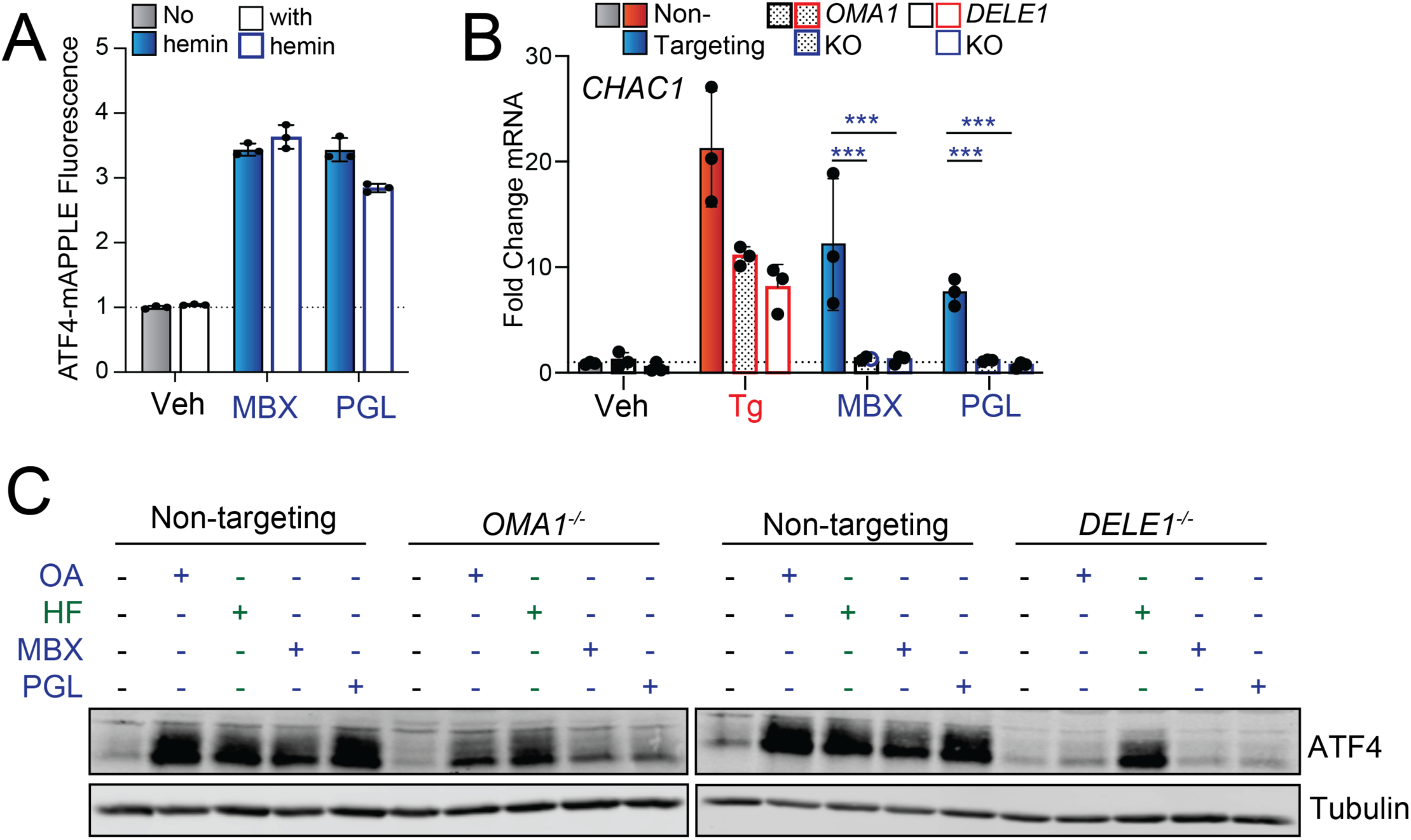
(Supplement to Figure 3). MBX-2982 and parogrelil activate the ISR through the OMA1-DELE1-HRI signaling axis. **A.** Fluorescence of the ATF4-mAPPLE ISR reporter, measured by flow cytometry, in HEK293T cells treated for 16 h with MBX-2982 (MBX; 10 µM), parogrelil (PGL; 10 µM), and/or hemin (10 µM), as indicated. **B.** Expression, measured by qPCR, of the ISR target gene *CHAC1* in non-targeting HEK293T cells or HEK293T cells CRISPRi-depleted *OMA1* or *DELE1* treated for 6 h with thapsigargin (Tg; 500 nM), MBX-2982 (MBX; 10 µM) or parogrelil (PGL; 10 µM). Error bars show SEM for n=3 replicates. ***p<0.005 for two-way ANOVA. **C**. Immunoblots of lysates prepared from non-targeting HEK293T cells or HEK293T cells CRISPRi-deleted of *OMA1* or *DELE1* treated for 6h with oligomycin A (OA; 500 nm), halofuginone (HF; 100 nM), MBX-2982 (MBX; 10 µM), or parogrelil (PGL; 10 µM).

**Figure S4.**
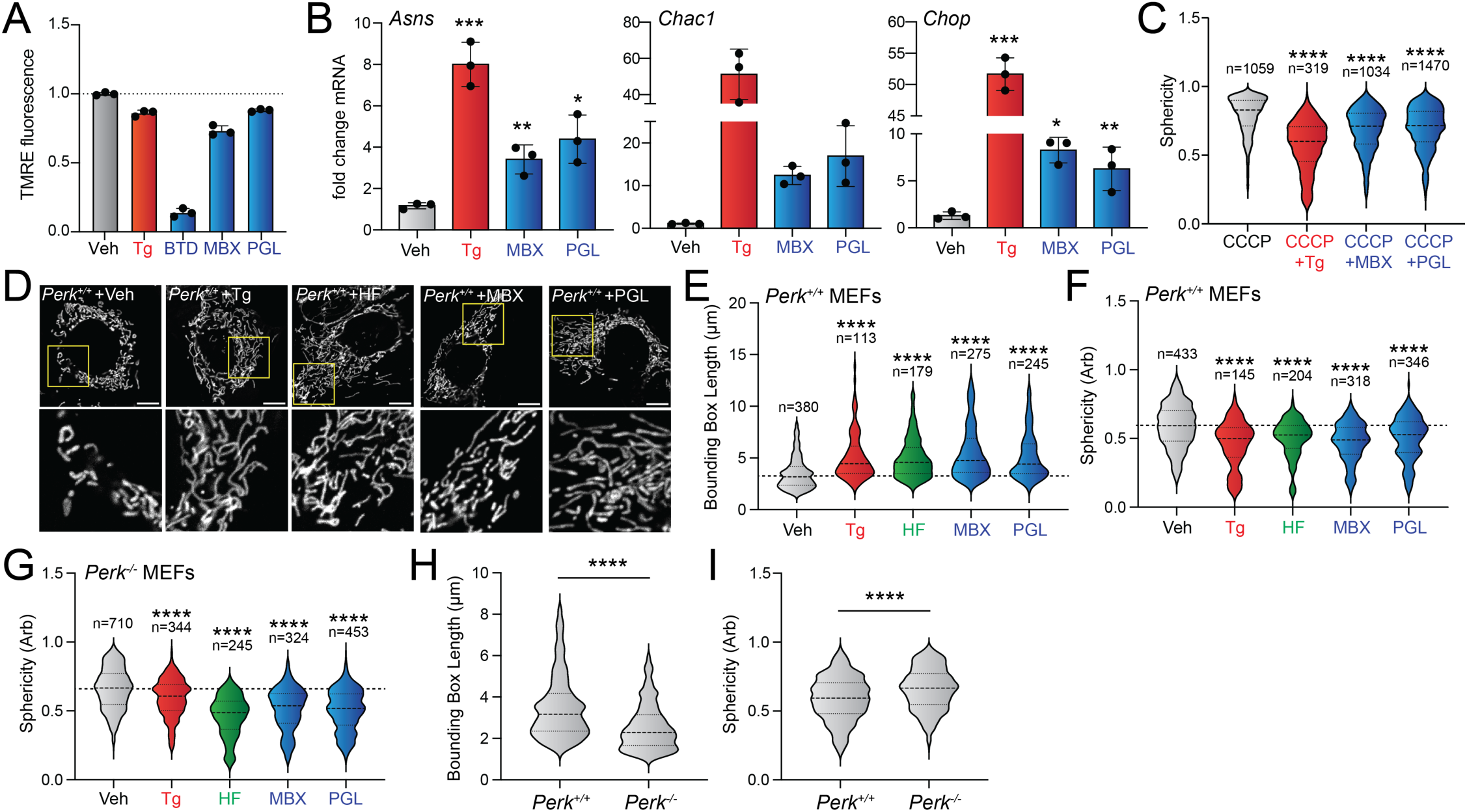
(Supplement to Figure 4) MBX and PGL promote protective mitochondrial elongation. **A.** TMRE fluorescence in HEK293T cells treated for 3 h with Thapsigargin (Tg; 500 nM), BtdCPU (BTD; 10 µM), MBX-2982 (MBX; 10 µM), Parogrelil (PGL; 10 µM). Error bar shows SEM for n = 3 replicates. **B.** Expression, measured by qPCR, of the ISR target genes *Asns*, *Chac1* and *Chop* in MEF cells treated for 6 h with thapsigargin (Tg; 500 nM), MBX-2982 (MBX; 10 µM) or parogrelil (PGL; 10 µM). Error bars show SEM for n=3 replicates. ***p<0.005 for two-way ANOVA. **C.** Quantification of sphericity from the entire dataset of images shown in **Fig 4D**. **D.** Representative images of TMRE stained *Perk^+/+^*MEF cells treated for 6 h with veh, thapsigargin (Tg; 500 nM), MBX-2982 (MBX; 10 µM), parogrelil (PGL; 10 µM). The inset shows a 3-fold magnification of the region shown by the yellow box. Scale bars 10 µm. **E, F.** Quantification of bounding box axis (E) and sphericity (F) from the entire dataset of images shown in D. **G.** Quantification of sphericity from the entire dataset of images shown in **Fig 4F**. The quantification of bounding box axis for *Perk^−/-^* MEFs treated with the indicated compound is shown in Fig. 4G. *p<0.05, **p<0.01, ***p<0.005, ****p<0.001 for one-way ANOVA (panels **A**,**B**), Kruskal-Wallis test (panels **F,G**), or Mann-Whitney test (panel **H,I**) relative to vehicle-treated cells.

**Figure S5.**
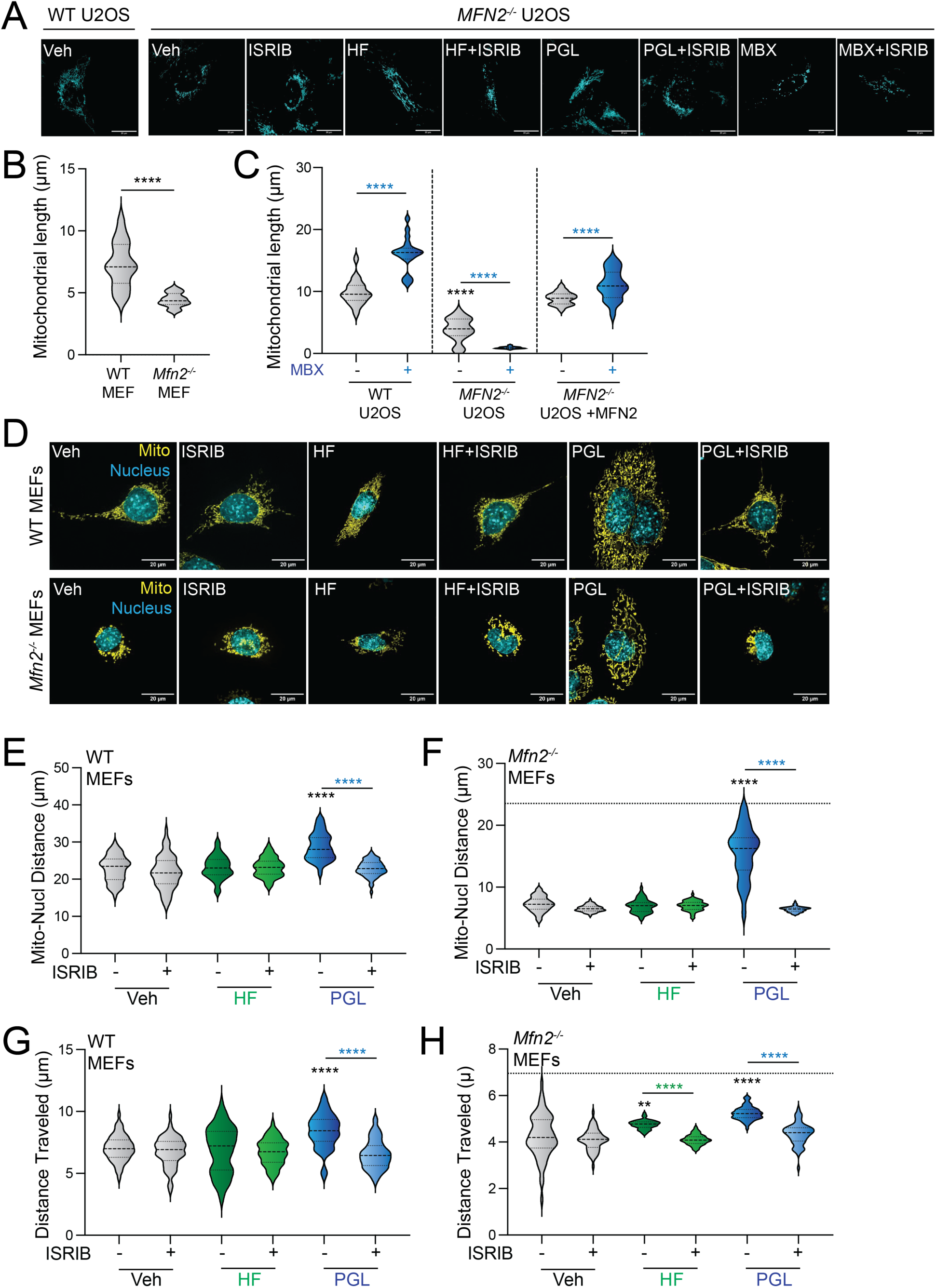
(Supplement to Figure 5). Pharmacologic ISR activation rescues mitochondrial morphology and motility in *MFN2*-deficient cells. **A**. Representative images of mitochondria in *Mfn2*-deficient MEFs treated for 6 h with halofuginone (HF; 100 nM); parogrelil (PGL; 10 µM), MBX-2982 (MBX; 10 µM), and/or ISRIB (200 nM) stained with TOMM20 antibodies (blue). **B**. Comparison of mitochondrial length in wild-type and *Mfn2*-deficient MEFs. **C**. Mitochondrial length in wild-type (WT), *MFN2*-deficient, and MFN2-deficient cells transfected with MFN2^WT^ U2OS cells treated for 6 h with MBX-2982 (MBX; 10 µM). **D**-**F**. Representative images (D) and quantification (E,F) of mitochondria-nuclear distance in wild-type MEF cells or *Mfn2-*deleted MEF cells treated for 6 h with halofuginone (HF; 100 nM); parogrelil (PGL; 10 µM), and/or ISRIB (200 nM) stained with TOMM20 antibodies (yellow) and Hoechst (blue). The dashed line in (F) shows mitochondria-nuclear distance in vehicle-treated wild-type cells for comparison. **G**,**H**. Mitochondrial distance traveled, measured by time lapse live cell imaging, in wild-type (left) or *Mfn2*-deficient WT cells (right) treated for 6 h with halofuginone (HF; 100 nM); parogrelil (PGL; 10 µM), and/or ISRIB (200 nM). The dashed line in (H) shows the distance traveled in vehicle-treated wild-type cells. Data shown were collected across two biological replicates. Violin plots show median and interquartile range. ****p<0.001 for Brown-Forsythe and Welch ANOVA. Black asterisks indicate comparison to vehicle-treated cells. Colored asterisks indicate comparison to cells co-treated with ISRIB.

**Figure S6.**
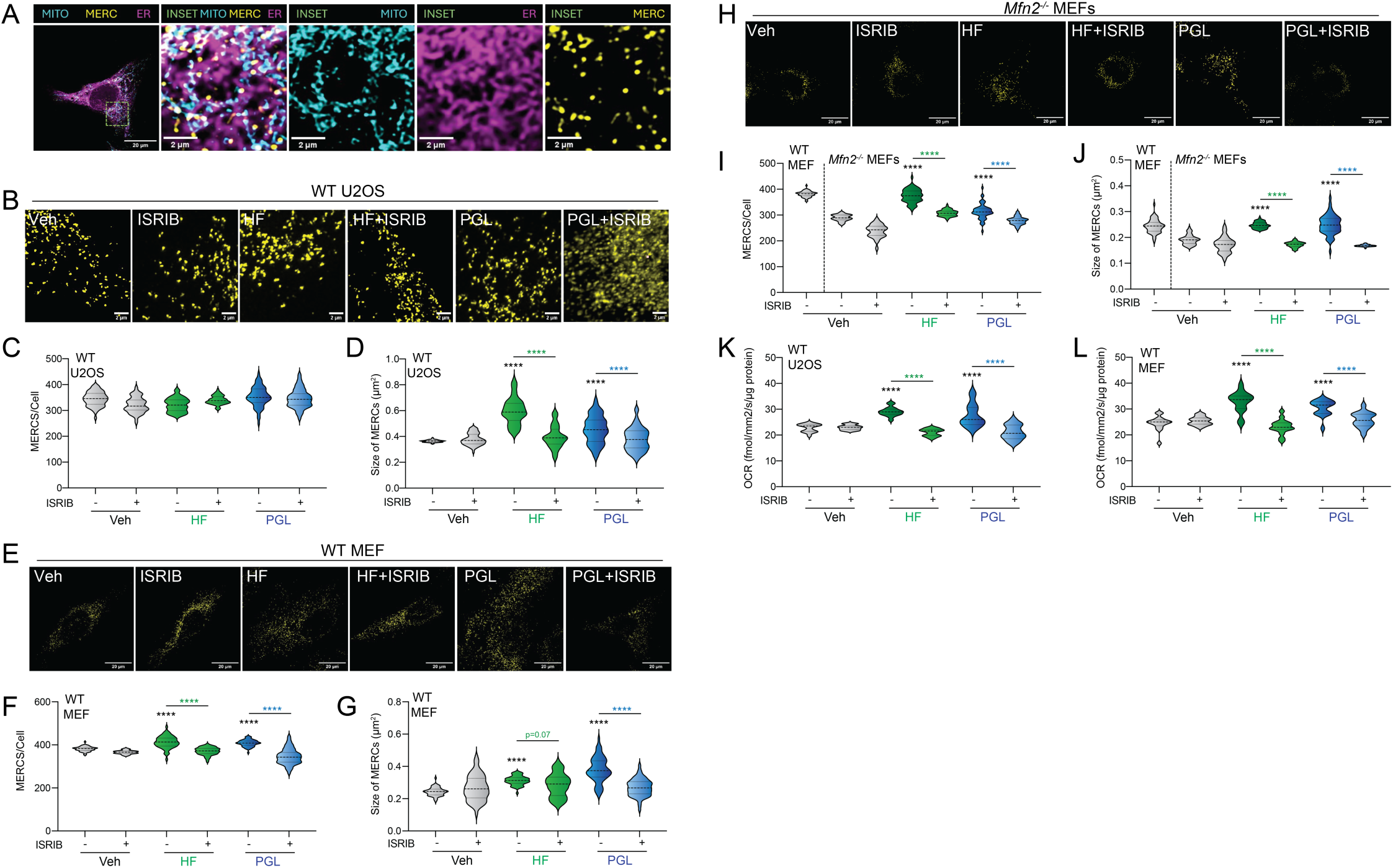
(Supplement to Figure 6). ISR activating compounds restore ER-mitochondrial contacts and enhance respiration in MFN2-deficient cells. **A**. Representative images of mitochondria stained with anti-TOMM20, endoplasmic reticulum (ER) stained with calnexin, and MERCs (PLA signal; see Fig. 1) in U2OS cells. **B-D**. Representative images (**B**) and quantification of MERC number (**C**) and size (**D**), measured by PLA, in wild-type U2OS cells treated for 6 h with halofuginone (100 nM), parogrelil (PGL; 10 µM), and/or ISRIB (200 nM). **E-G**. Representative images (**E**) and quantification of MERC number (**F**) and size (**G**), measured by PLA, in wild-type MEF cells treated for 6 h with halofuginone (100 nM), parogrelil (PGL; 10 µM), and/or ISRIB (200 nM). **H-J.** Representative images (**H**) and quantification of MERC number (**I**) and size (**J**), measured by PLA, in *Mfn2*-deficint MEF cells treated for 6 h with halofuginone (100 nM), parogrelil (PGL; 10 µM), and/or ISRIB (200 nM). PLA signal in wild-type MEF cells is shown for comparison. **K,L.** Oxygen consumption rate (OCR), measured using the Resipher system, in wild-type U2OS (**K**) and MEF (**L**) cells treated for 6 h with halofuginone (HF; 100 nM), parogrelil (PGL, 10 µM), and/or ISRIB (200 nM). ****p<0.005 for one-way ANOVA. Black asterisks show comparison with vehicle-treated cells and colored asterisks represents comparison with ISRIB co-treatment.

